# Role of DNA-DNA sliding friction and non-equilibrium dynamics in viral genome ejection and packaging

**DOI:** 10.1101/2023.04.03.535472

**Authors:** Mounir Fizari, Nicholas Keller, Paul J. Jardine, Douglas E. Smith

## Abstract

Many viruses eject their DNA via a nanochannel in the viral shell, driven by internal forces arising from the high-density genome packing. The speed of DNA exit is controlled by friction forces that limit the molecular mobility, but the nature of this friction is unknown. We introduce a method to probe the mobility of the tightly confined DNA by measuring DNA exit from phage phi29 capsids with optical tweezers. We measure extremely low initial exit velocity, a regime of exponentially increasing velocity, stochastic pausing that dominates the kinetics, and large dynamic heterogeneity. Measurements with variable applied force provide evidence that the initial velocity is controlled by DNA-DNA sliding friction, consistent with a Frenkel-Kontorova model for nanoscale friction. We confirm several aspects of the ejection dynamics predicted by theoretical models. Features of the pausing suggest it is connected to the phenomenon of “clogging” in soft-matter systems. Our results provide evidence that DNA-DNA friction and clogging control the DNA exit dynamics, but that this friction does not significantly affect DNA packaging.

## Introduction

Many double-stranded DNA viruses, including tailed bacterial viruses (“phages”) and some human/animal viruses such as herpesviruses, employ ATP-powered molecular motors to package DNA into preformed viral capsid shells (1,2,3). The DNA is very tightly packed, reaching densities as high as ∼660 mg/ml (∼50% volume fraction). This density is well above that at which short DNA segments in solution form liquid crystal phases. X-ray scattering measurements indicate that the spacing between adjacent ∼2 nm diameter DNA segments is only ∼0.5-1 nm. The remaining volume is occupied by thin layers of water molecules and ions (4-6).

Viral packaging motors exert large forces to translocate DNA into the procapsid through a portal nanochannel against large forces resisting DNA confinement that arise from factors including electrostatic self-repulsion of DNA segments, DNA bending rigidity, and changes in DNA hydration and entropy (7-10). Part of the work done by the motor is stored as potential energy, giving rise to what has been termed “DNA pressure” or “internal force”. While this force resists packaging, it plays an important role in driving DNA ejection when the virus infects a host cell (11-15).

The dynamics of motor-driven DNA packaging in phages phi29, lambda, and T4 have been extensively studied via single-molecule measurements with optical tweezers (3, 16-24). The motors generate forces >60 pN, placing them among the strongest known ATP-powered motors. DNA packaging begins rapidly, at rates ranging from ∼150-2000 bp/s depending on virus and condition, but then slows as the capsid fills (16-18, 23). This is due, in part, to the buildup of internal force that loads the motor (20,24). In phage phi29, the force reaches ∼25 pN at the end of packaging (20,24,25), which is consistent with theoretical predictions and the experimentally determined DNA ejection force in phage lambda, which packs DNA to similarly high density (25,26).

DNA ejection from phages T5 and lambda has been measured *in vitro* with fluorescent labeled DNA (12, 29-31). Although the force that drives ejection is highest at the beginning of the process and decreases as ejection proceeds, a notable finding was that the DNA ejection velocity has an initially increasing trend. It was proposed that this is due to hydrodynamic drag forces acting on the confined DNA that decrease with decreasing DNA packing density. After about half of the genome is ejected there is another regime in which the velocity decreases, which is attributed to decreasing internal force. Viral DNA ejection *in vivo* has also been studied (14,15,32) and is more complex because it involves interactions with the cytoplasm or nucleoplasm. We focus here on *in vitro* ejection to investigate the fundamental question of how DNA mobility is affected by the very high-density packing. This question is also of interest in soft-matter/polymer physics since DNA is a model for semiflexible polymers in general and viral packaging is a model for studying how confinement affects polymer dynamics.

Another surprising finding is that long pauses of up to ∼100 s were observed during T5 DNA ejection (12, 31), but no pauses were observed with phage lambda. The T5 studies also found that the ejection dynamics depend on the concentration of dye used to label the DNA (33), which is likely due to the dye binding affecting the physical properties of DNA. It is thus advantageous to develop methods to measure ejection without dye labeling (34).

Viral DNA ejection has also been considered in many theoretical studies, including both analytical theories and dynamic simulations (35-51). One early model proposed that ejection is slowed due to hydrodynamic drag forces, such that the velocity, *v* = *µF*, is proportional to the driving force *F*, where µis a mobility coefficient. A reptation-type polymer model was used to predict that µincreases as ejection proceeds due to decreasing DNA confinement (35). This model was used in interpreting the lambda ejection results (30). However, it has been proposed by other researchers that sliding friction between closely packed DNA segments could be the main source of friction rather than hydrodynamic drag (36,37,52).

We introduce a method to probe the DNA mobility by using optical tweezers to pull single DNA molecules out from single phi29 viral capsids. The method provides higher resolution measurement of DNA exit dynamics, avoids the use of fluorescent dyes, and allows us to study how exit velocity depends on driving force. We find that during the initial stages of DNA exit the velocities are much lower than previously reported for T5 and lambda, and there is a much sharper increase in the velocity as ejection proceeds. We find evidence that DNA-DNA sliding friction controls the initial ejection speed rather than hydrodynamic drag. We also observe frequent pauses during ejection but find that they have a different character than those observed in the phage T5 studies by fluorescence imaging.

## Materials and Methods

Methods for initiating and measuring DNA packaging are similar to those described previously (53). Fiberless phage phi29 proheads (empty capsids) and recombinant gp16 packaging ATPase proteins were produced as described previously (54,55). A biotin labeled 25-kbp dsDNA packaging substrate was produced via PCR as described previously (56).

Prohead-motor complexes were prepared by mixing 2 µg of proheads with 0.4 µg of gp16 ATPase in 10 µL 0.5x TMS buffer (25 mM Tris·HCl, pH 7.5, 50 mM NaCl, 5 mM MgCl_2_) and incubating for 4 min, before adding a final concentration of 0.5 γS-ATP to a final concentration of 0.5 mM. Complexes were then incubated at 4°C for 1 hr before use. The DNA constructs were tethered to streptavidin-coated 2.3 μm microspheres and the prohead-motor complexes were attached to 2.1 μm protein G-coated microspheres coated with IgG antibodies against the phage capsid protein, gp8.

Packaging was initiated in a fluid chamber in the optical tweezers instrument by bringing a microsphere carrying DNA into near contact with a microsphere carrying prohead-motor complexes in a buffer containing 0.5 mM ATP, as described previously (53). In measurements where Na^+^ was the dominant ion screening the DNA charge, the packaging buffer contained 25 mM Tris·HCl, pH 7.5, 50 mM NaCl, 5 mM MgCl_2_, whereas measurements with Mg^2+^ as the dominant screening ion were conducted in a buffer containing 1 mM Tris·HCl pH 7.5 and 30 mM MgCl_2_. After packaging close to 100% of the genome length (which takes, on average, ∼8 minutes), complexes were moved into a region of the fluid chamber containing an identical buffer but without ATP. A gentle flow was maintained so that no ATP could diffuse into this region. Based on the measured kinetic rates of the packaging motor, any bound ATP would be spontaneously hydrolyzed on a timescale less than the time it takes to finish the solution exchange and initiate measurement of the DNA exit (i.e., in <1 s) (20, 57).

Measurements were made with a dual-trap optical tweezers system at room temperature (∼23 °C) as described previously (53). The force signal was recorded at 1 kHz and measurements were conducted in force-clamp mode, where the position of the movable trap is adjusted by a feedback control system to keep the applied force on the DNA constant. The tweezers were calibrated as described previously (58). The tether length was computed from the measured force vs. fractional extension relationship which is accurately described by the extensible worm-like chain model for DNA elasticity. The instantaneous capsid filling level was computed by subtracting the current tether length from the initial tether length at the start of packaging. DNA exit velocities were calculated by linear fits to DNA tether length vs time in a 0.5 s sliding window. These velocities were averaged together in 5% filling bins to determine a velocity vs filling profile for each complex, and then averaged across complexes to calculate the mean velocity vs. filling level. DNA exit from an individual complex was required to cover at least half of a filling bin to contribute to the average. Pauses were detected using the method described by Kalafut and Visscher (59). A lower resolution limit for both pause duration and change in tether length between pauses was determined by measuring experimental noise/drift levels in control datasets (static DNA tethers held at constant force) and comparing the velocity in a time interval Δt during the pause or between pauses to 1 standard deviation from the mean of the distribution of velocities of control data measured in time intervals of duration Δt.

Internal force values were determined based on previous optical tweezer measurements via interpolation of the mean force-filling relationship (24,27). Inferred internal force and the applied pulling force were added for each complex then averaged across different complexes for each filling bin. Error bars on averaged quantities represent standard errors calculated by bootstrap sampling.

## Results

### Measurement of DNA exit trajectories

The experimental method is illustrated in Figure 1A and builds on methods we developed previously for measuring motor-driven packaging (28). We first initiate packaging by bringing a trapped microsphere with tethered DNA near a second trapped microsphere carrying capsid-motor complexes in a solution with ATP. Packaging proceeds until the capsid filling (% of the 19.3 kbp viral genome length in the capsid) reaches ∼100% and then the complex is moved to a region without ATP. This causes the motor to release its grip and the DNA exits the capsid. We apply a small 5 pN force to keep the DNA stretched so the length that has exited can be measured. Further details are given in the methods section.

**Figure 1.**
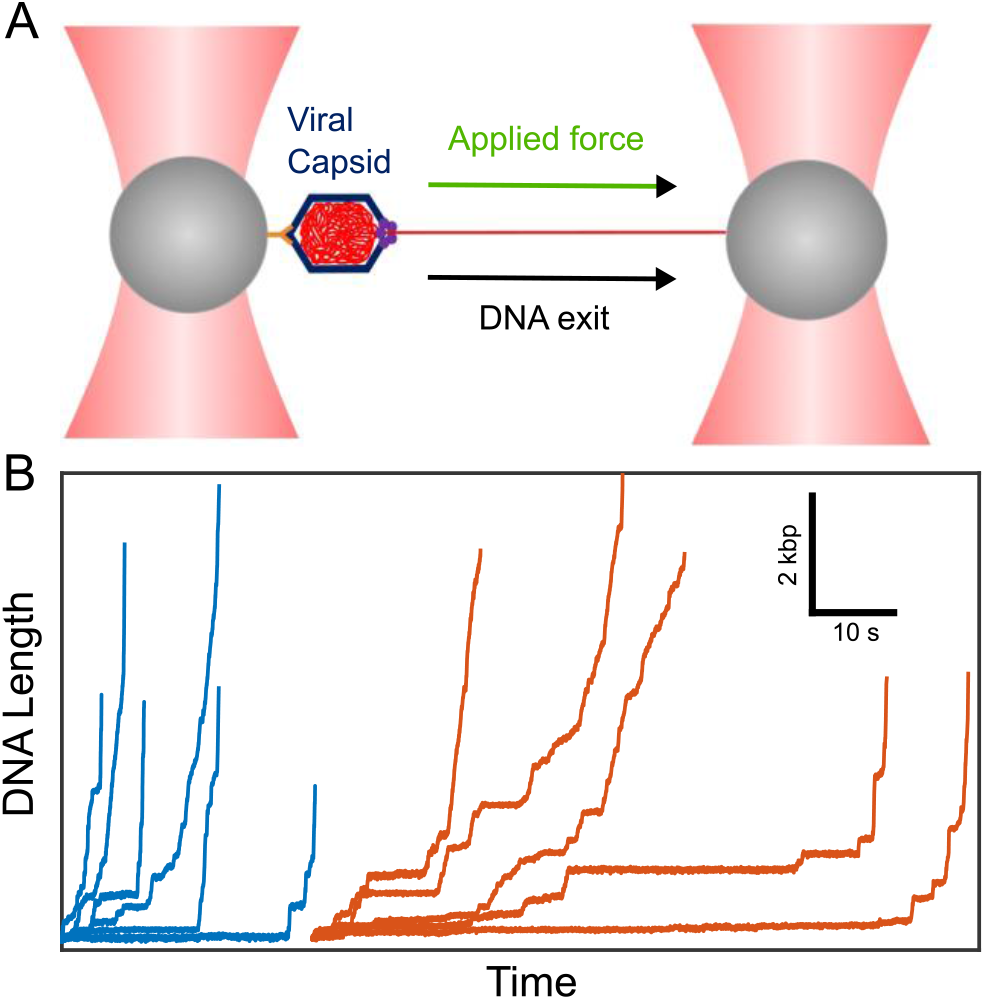
(A) To probe the mobility of the tightly packed DNA in single phage phi29 viral capsids we pull the DNA out through the portal nanochannel using optical tweezers. The capsid is attached to a microsphere held in one optical trap (left) and the end of the DNA is attached to a second microsphere held in a second trap (right). (B) Measurements of the length of DNA that has exited vs. time with a 5 pN pulling force and with Na^+^ the dominant ion screening the DNA charge (blue plots) or with Mg^2+^ (red plots). Each line is a measurement on a single complex, starting with ∼100% of the genome length packaged and ending when the velocity exceeds the maximum measurable with the setup (∼12 kbp/s).

Examples of measured DNA exit trajectories are shown in Figure 1B. Striking features are that the initial velocities are extremely low (much lower than observed in the prior T5 and lambda studies with dye-labeled DNA) and there are pauses of widely varying duration across the whole range of measured capsid filling levels. As DNA exit proceeds, the pausing decreases and the velocity increases, eventually exceeding ∼12 kbp/s, which is the maximum we can measure with our instrument. At this point, the velocity is so high that it has negligible influence on the overall DNA exit time.

We measured an ensemble of events to determine average DNA exit velocity and assess dynamic heterogeneity. Transient velocity versus filling level was determined and averaged across complexes in 5% bins. A trend of increasing average velocity with decreasing filling level is observed regardless of whether the pauses are included or excluded in the velocity calculations (Figure 2A and Supplementary Figure S1). We hereafter use the term “velocity” to refer to transient velocity not including pauses. Strikingly, the average velocity increases exponentially as the capsid filling level decreases from ∼100-80%, going from ∼200 bp/s to ∼2,000 bp/s. In Figure 2 we plot average velocities from 80-100% filling where the transient velocities for all complexes were low enough to be measured. This range dominates the overall ejection time. Many individual complexes could also be measured down to much lower filling levels (Supplementary Figure S2A).

**Figure 2.**
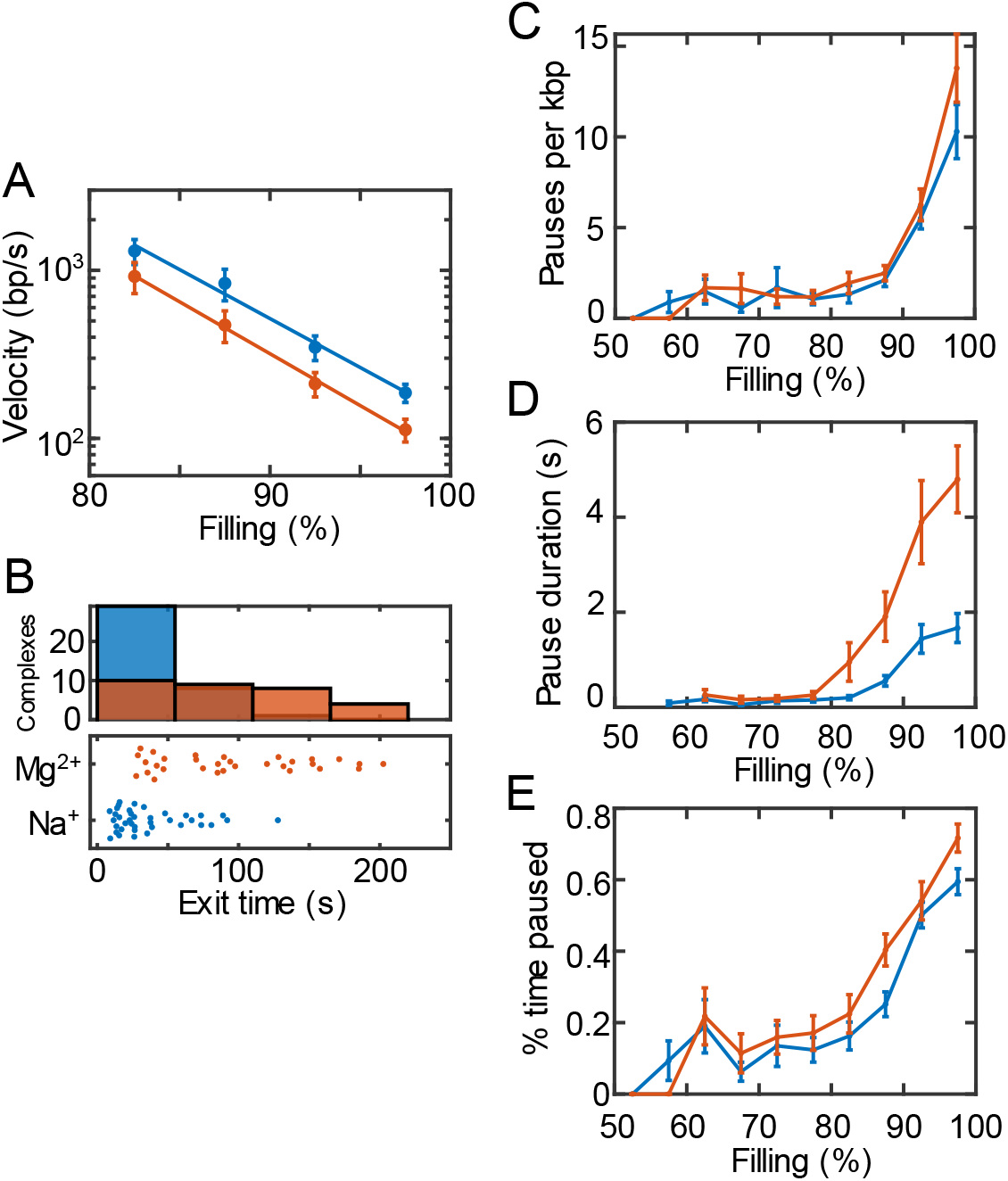
(A) Average DNA exit velocity, not including pauses, vs. capsid filling level in the high packing density regime in the Na^+^ screening condition (blue, n = 39 events) or Mg^2+^ screening condition (red, n = 31). Measurements on individual complexes extending to lower filling levels are presented in the Supplementary Figure S2. (B) Distributions of overall DNA exit times measured for individual complexes (top) and corresponding measured values for individual complexes (bottom). (C) Average durations and (D) average frequency of pauses versus capsid filling levels. (E) Fraction of time DNA exit is paused vs. capsid filling. Panels B-E use the same color code as panel A.

### Pausing during DNA exit and nonequilibrium dynamics

We observe significant pausing during DNA exit. The pauses occur stochastically across all capsid filling levels down to ∼50% and have widely varying durations as long as ∼15 s. We measure individual pauses with high resolution and determine their frequencies and durations. The mean frequency and mean duration decrease monotonically with decreasing capsid filling level (Figure 2C and 2D). The percent time spent paused decreases from ∼60% near 100% filling to ∼20% at 80% filling (Figure 2E). These pauses dominate the overall ejection kinetics, comprising 57% (± 3%) of total exit time.

We also observe broad heterogeneity in DNA exit dynamics between different individual complexes, clearly seen in the length vs. time and velocity measurements (Figure 1B and Supplementary Figure S2A). Whereas the trend of increasing velocity with decreasing filling level is obeyed on average, exit velocities in individual complexes also exhibit large fluctuations in time. There are even instances where the velocity transiently decreases as the filling level decreases (Supplementary Figure S3). The velocity limit of 12,000 bp/s is reached at quite variable filling levels in each complex, ranging from ∼40-80% filling (mean of 73 ± 3%). Overall exit times for different complexes vary widely, ranging from ∼10-100 s (Figure 2B), consistent with the process being controlled by nonequilibrium dynamics of the tightly packed DNA.

### Dependence on ionic screening

Ionic screening of the DNA phosphate backbone charge is expected to be an important parameter because it influences the strength of electrostatic DNA-DNA interactions (60,61). Phage lambda ejection studies with fluorescent DNA found that increasing the screening, thus reducing DNA charge repulsion, slowed ejection (29,30). We repeated our measurements with higher screening by omitting NaCl and increasing MgCl_2_ to 30 mM; divalent cations result in an ∼25% lower effective DNA charge per unit length (62). DNA-DNA interactions are net repulsive in both conditions, but weaker with Mg^2+^ (63,64). We find that the mean DNA exit velocity is reduced (Figure 2A). Velocities measured for individual complexes are shown in Supplementary Figure S2B, again showing large heterogeneity. Whereas the trend of exponentially increasing average exit velocity remains similar, velocities are reduced by approximately 50% across the whole range compared to the lower screening condition. Both the mean pause duration and mean pause frequency follow a similar trend with filling as in the lower screening condition (Figure 2C&D). Pause duration is increased significantly in the high screening condition and pause frequency increases at the highest filling levels, leading to a ∼30% increase in the fraction of time spent paused during DNA exit. As shown in Figure 2B, the distribution of DNA exit times shifts towards higher values.

### Dependence on driving force

A unique advantage of our method using optical tweezers is that we can apply controlled pulling forces to vary the force driving DNA exit. An unanswered question in the field is whether frictional forces restricting the movement of the tightly packed DNA are predominantly due to hydrodynamic drag or some type of DNA-DNA sliding friction. For hydrodynamic drag, a linear relationship between velocity and force, *v* = *µF*, was predicted, where *µ* is a mobility constant. This relationship was used to interpret the phage lambda ejection data, but this model has not been tested experimentally. To test this, we repeated DNA exit measurements with a significantly higher applied pulling force of ∼20 pN (Supplementary Figure S4).

We observe qualitatively similar exit dynamics as with 5 pN pulling force, but the mean velocity is higher across the whole range of capsid fillings (Figure 3A). Velocities measured for individual complexes are shown in Supplementary Figure S2C, again showing significant heterogeneity, and the frequency and average duration of pauses is decreased (Figure 3B and 3D). A notable finding is that the relationship between exit velocity and total driving force (sum of the internal force and applied pulling force (20,24)) is not linear as predicted by hydrodynamic drag models. This is shown in Figure 3C where the ratios of the exit velocities and ratios of total driving forces with the high and low applied forces are compared. The magnitude of the friction force opposing DNA exit is equal to the total driving force since the molecular motion is overdamped on the timescale of measurement. The ratio of the velocities is notably higher than the ratio of forces across the measured range of filling levels (Figure 3C), which shows that the relationship *v* = µ*F* does not hold.

**Figure 3.**
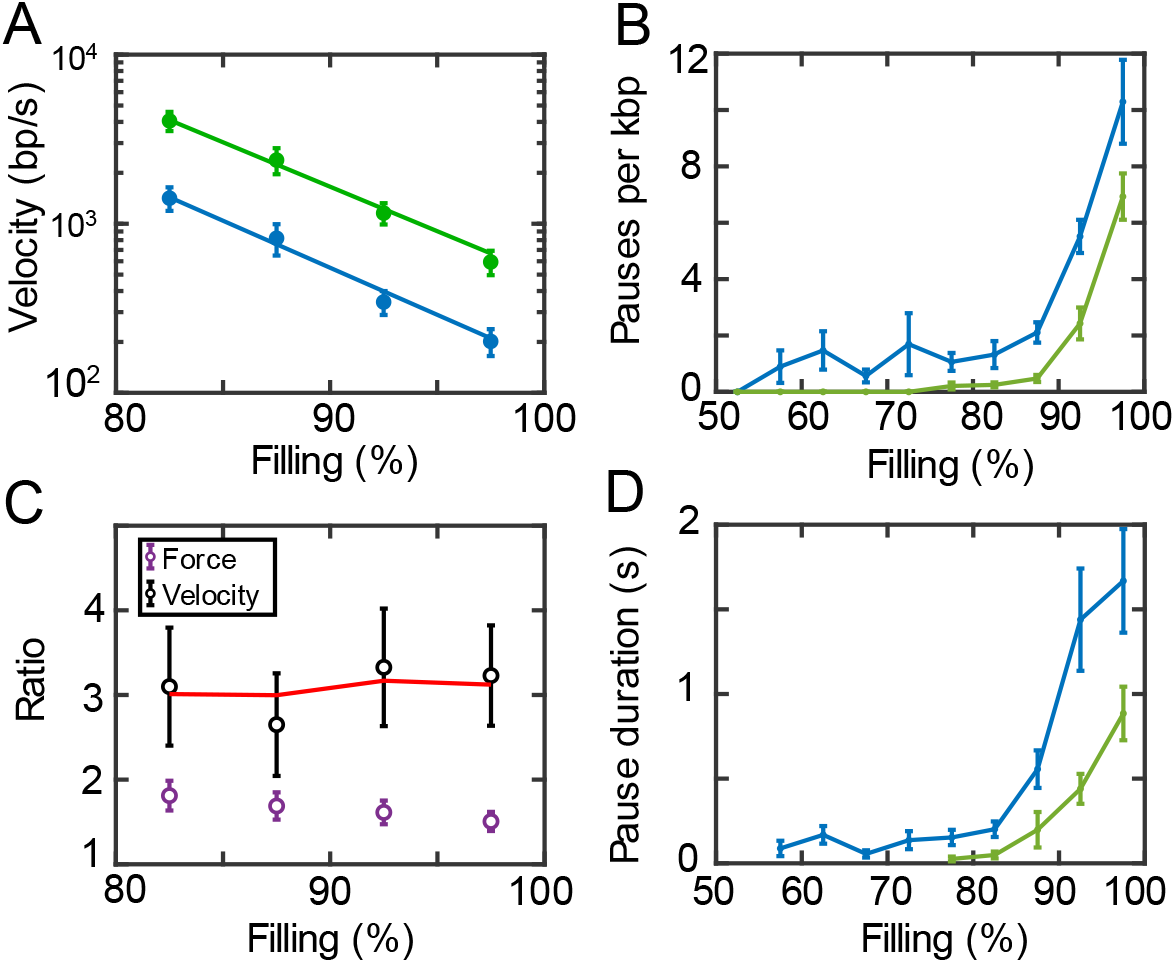
(A) Average DNA exit velocity, not including pauses, vs. capsid filling with 5 pN (blue) and 20 pN (green) applied force. Measurements on individual complexes extending to lower filling levels are presented in the Supplementary Figure S2. (B) Ratio of average velocities (black) compared with ratio of total driving forces (purple) with the 20 pN and 5 pN applied forces, versus capsid filling. The ratios of velocities predicted by the Prandtl-Tomlinson relationship with d = 0.34 nm is shown by the red line. (C) Average pause frequency, and (D) average pause duration vs. capsid filling (with Na^+^ screening) with 5 pN (blue, n=39 events) or 20 pN applied force (green, n=42).

## Discussion

DNA ejection is driven by the internal forces resisting tight confinement of the charged biopolymer that build up during packaging. The dynamics of ejection, however, depends on friction forces that resist DNA exit. The nature of this friction has been considered in some theoretical models, but experimental data has not been available to investigate this.

In one influential early model, Gabashvili and Grosberg considered that the main source of the friction could be hydrodynamic drag acting on the DNA inside the capsid (35). They proposed that, due to the high density packing, DNA segments might undergo a reptation-like motion during ejection, akin to that of entangled polymers in concentrated solutions. Entanglements restrict a polymer to a move primarily in a fluid-filled tube-like volume parallel to its own contour. The viral DNA is envisioned to behave as a self-entangled polymer and to experience hydrodynamic drag force proportional to exit velocity *v*, such that *F* = *v*/*µ*, where *µ* is a mobility factor. Applying classical continuum fluid mechanics, µwas predicted to increase as the DNA packing density decreases because the confining tube diameter would decrease, resulting in increased hydrodynamic shear forces (35). This model could thus explain a trend of initially increasing DNA exit velocity (provided that the internal force does not decrease too rapidly as the DNA ejects). However, the assumption that the friction force at each capsid filling level is proportional to DNA exit velocity is inconsistent with our findings (Figure 3C). Moreover, the predicted hydrodynamic drag forces are much smaller than those we detect. For example, when ejecting at 93% capsid filling, we detect a friction force of ∼20 pN at a mean exit velocity of ∼300 bp/s, which is much higher than the ∼0.01 pN predicted by the hydrodynamic model for this velocity (see the Supplementary text).

Other researchers have proposed that sliding friction between adjacent, tightly packed DNA segments may be important. Odijk considered the possibility of simple Coulomb (macroscopic solid-like) sliding friction (36). The friction force depends on normal forces that press adjacent segments together, proposed to be proportional to internal force. Since these internal forces decrease as ejection proceeds, this model could also explain a trend of increasing velocity as ejection proceeds. A similar idea was explored by Ghosal (37), who modeled DNA as a semiflexible elastic rod initially arranged in an inverse spool and subject to Coulomb sliding friction. However, the friction force assumed in these models is, at each capsid filling level, independent of DNA exit velocity, which is inconsistent with our findings (Figure 3C).

Rather, our measurements provide evidence that DNA exit is controlled by a type of nanoscale biopolymer sliding friction described recently. Ward *et al*. used macromolecular crowding to press two actin filaments together and measured the force required to slide one past the other (65). The friction force was found to increase logarithmically with velocity. This finding was shown to agree with the predictions of a Frenkel-Kontorova-type model for nanoscale friction, that accounts for effects of the periodic atomic-scale “bumpiness” of individual polymers and the role of thermal fluctuations. The model predicts that friction force follows *F* =(*kT*/*d*)*log*(*v*/µ), known as the Prandtl-Tomlinson relationship, where *µ* is an effective mobility factor, *kT* the thermal energy, and *d* the spacing between monomers in the biopolymer (65). This was shown to fit the actin data with *d* = 5 nm set equal to the monomer spacing for F-actin filaments.

Our measurements provide evidence that an analogous type of DNA-DNA sliding friction acts during the initial stages of slow DNA exit. The ratios of exit velocities we measure with high and low externally applied forces agree with the predictions of the Prandtl-Tomlinson relationship with *d* set to the 0.34 monomer spacing (per bp) of dsDNA (Figure 3C). This agreement holds over the whole range of filling levels where we measure exponentially increasing exit velocity. This friction likely contributes to the very long timescale for DNA reorganization we observed in previous studies in which packaging complexes were stalled at ∼75% filling level (28). As ejection proceeds to lower filling levels and the average spacing between DNA segments increases, a crossover to a regime where the friction is dominated by hydrodynamic drag is expected (29, 30, 63). However, as discussed above, at this point the exit velocity is so fast that it has a negligible effect on the overall kinetics.

Since systems with sliding friction sometimes exhibit “slip-stick” dynamics (66, 67), we considered whether such behavior could explain the pausing we observe during DNA exit. Within the Frenkel-Kontorova model, stick-slip dynamics can occur if the interaction potential is sufficiently large compared to the thermal energy and work done by driving forces. The distance between pauses during ejection would be predicted to be on the scale of the periodicity of the interaction potential, or ∼0.34 nm for DNA, and the pause durations exponentially distributed. The pauses we observe, however, have much larger distances between them (∼10-100 bp) (Figure 4A) and their durations are not exponentially distributed (Figure 4B). Thus, our results do not support slip-stick friction as a mechanism for pausing.

**Figure 4:**
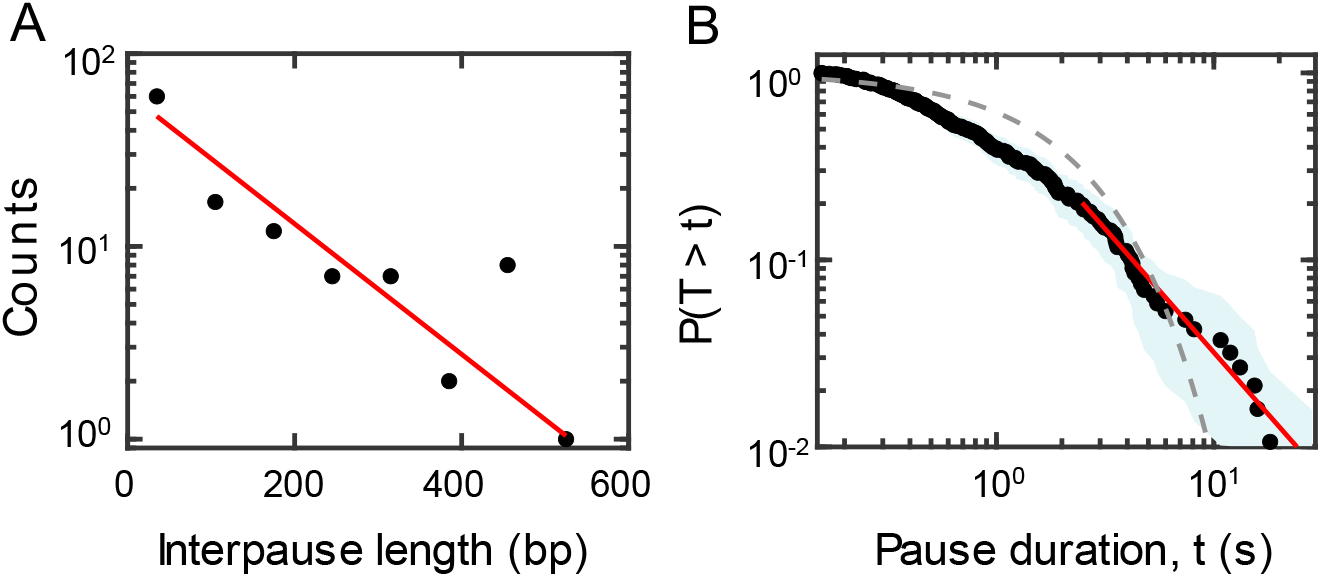
(A) Histogram of the lengths of DNA released between each pause. Line shows a fit to an exponential distribution. (B) Complementary cumulative distribution function (i.e., the probability of finding a pause with duration T that is greater than a given value t) at the 90-95% filling level across all complexes in the Na^+^ screening condition (black points). The blue band shows the 95% confidence interval. Dashed lines show an exponential fit to the distribution (gray) and a power law fit to the distribution tail determined by the Clauset-Shalizi-Newman method (82) (red).

Our findings further allow us to consider whether DNA-DNA sliding friction has any significant effect on motor-driven DNA packaging. Since the friction is mainly dependent on the spacing between adjacent DNA strands, we expect that at equivalent capsid filling levels the nature of the friction will be similar during ejection and packaging. However, since packaging is much slower than ejection, the Prandtl-Tomlinson relationship implies much lower friction forces during packaging. Our measurements find that the average DNA translocation velocity during packaging, at room temperature and with saturating ATP, decreases from 55 bp/s to 11 bp/s as the filling increases from 80% to 100%. At these velocities the Prandtl-Tomlinson relationship predicts friction forces <0.1 pN (see Supplementary text), which are negligible compared to the internal forces resisting packaging (∼5-30 pN).

We note that if the phi29 motor were translocating at the maximum value of ∼160 bp/s and did not slow with capsid filling as observed (16,20,24), then the predicted friction forces would be significant (∼25 pN). One of the reasons the motor evolved to down-regulate DNA translocation velocity with increasing filling level may be to minimize friction forces that would otherwise result in an additional energy cost during packaging and possible motor stalling. On the other hand, the lambda and T4 motors translocate considerably faster than phi29 (up to ∼10x faster in the case of T4) (17,18), so it is possible that DNA sliding friction could play a role in those systems.

DNA ejections from phages T5 and lambda, which pack DNA to similar density as phi29, were studied using fluorescent labeled DNA. Mangenot *et al*. developed a method in which T5 phages were attached to a surface, ejection was triggered with purified host cell membrane protein, and the exiting DNA was stretched in a flow and imaged by fluorescence microscopy (12). More detailed studies were conducted by Chiaruttini *et al*. (31), and the method was extended to study phage lambda by Grayson *et al*. and Wu *et al*. (29,30).

For phage lambda, Grayson *et al*. reported initial average ejection velocities of ∼25,000 bp/s increasing to ∼30,000 bp/s at 80% filling (29). Using a lower dye concentration, Wu *et al*. measured ∼3,000 bp/s at 85% filling, rising to ∼6,000 bp/s at 80% filling (30). In both cases the velocity initially increased approximately linearly with decreasing filling level. In contrast, in a similar ionic condition, for phi29 with unlabeled DNA we measure a much lower initial average velocity of ∼200 bp/s at ∼100% filling and a much sharper, exponential increase in the velocity to ∼2000 bp/s at 80% filling. Wu *et al*. also studied the dependence of lambda ejection on ionic screening and observed that it slows with increased screening, attributed to decreased internal force due to reduced electrostatic DNA-DNA repulsion (30). Although our velocities are systematically lower and dependence on filling level is different, we observed a similar trend of decreased velocity with increased ionic screening.

For phage T5 Chiaruttini *et al*. reported much higher DNA exit velocities than we observe for phi29 with unlabeled DNA: ∼7,000 bp/s at 100% filling (31), which is ∼35-fold higher than we measure in a similar ionic condition. Their velocity figure also averaged in pauses, which implies even higher transient velocities. They also observed a less than linear increase in velocity from 100% to 80% filling, compared to the much sharper (exponential) increase we observe.

We observe frequent pausing that dominates the overall DNA exit kinetics. While pausing was also observed in the T5 studies (12,31,33), the character of the pausing we observe is quite different. The pauses we observe occur stochastically with a frequency and duration that decreases with decreasing filling level, whereas those reported for T5 occurred mainly around a few specific filling levels (94 ± 1%, 88 ± 4%, and 60 ± 12%). It was suggested that those pauses might be caused by liquid-crystal phase transitions of the DNA, a proposal guided by electron microscopy studies of partly ejected T5 phages which found evidence for partial ordering of the DNA in liquid-crystal-like phases (33). However, it is unclear if this explains the pauses since they did not occur at the same density levels as the phase transitions.

The large differences in DNA exit dynamics we measure compared with those reported in the T5 and lambda studies is surprising since these phages all pack DNA to similarly high densities. Recent X-ray scattering studies that probe the spacings between packed DNA segments in phage particles suggest that the packing density is only slightly higher in phi29 (615 ± 50 mg/ml vs. 540 ± 20 mg/ml for T5 and 525 ± 20 mg/ml for lambda), where the differences are not much larger than the uncertainties in the measurements (68). Other differences are that lambda and T5 have genomes and capsid sizes that are ∼2.5-fold and ∼6-fold larger than phi29, respectively, and have isometric (spherically symmetric) capsids whereas phi29 is prolate (slightly elongated). These differences would make the DNA bending energy per unit length higher for phi29, which would increase the internal force. However, this difference would be expected to cause the DNA exit velocity to be higher rather than lower.

Another difference is that our method does not require the DNA to be labeled with dye. Studies with phage T5 found that the ejection dynamics is affected by the concentration of dye used to label the DNA (33). This likely occurs due to changes in the physical properties of DNA such as contour length, bending rigidity, and net charge caused by dye binding (69,70). The studies of sliding friction between actin filaments also showed that the sliding friction is sensitive to perturbations to the biopolymer structure (65). Changes in DNA structure caused by dye binding might therefore affect ejection dynamics by altering DNA-DNA sliding friction.

There have been a wide variety of theoretical studies of DNA ejection: we identified 48 published studies, including analytic models and simulations (see Supplementary references for a full list). Above we discussed three analytic models that proposed explicit models for friction during ejection but showed that these are inconsistent with our findings. Most of the other analytic studies have focused on predicting energetics of ejection and/or scaling laws for ejection time. Several considered ejection dynamics but predict only monotonically decreasing velocity, attributed to decreasing driving force as ejection proceeds (44,48-51). Only one of these studies, by Wang *et al*. (51), predicts an initially increasing velocity. This study modifies the hydrodynamic model (35) by adding a compressive osmotic force that increases drag force per length of the confined DNA. However, it again predicts a much smaller friction force (∼0.4 pN) than we measure. Notably, all of these models assume sliding friction between adjacent DNA strands is negligible compared to hydrodynamic forces.

Many simulation studies have also been conducted, all using coarse-grained bead-spring polymer DNA models. Unlike in analytic models where DNA-DNA sliding friction would need to be explicitly included, simulations have the potential to account for such friction via steric interactions between beads. However, it has remained unclear to what extent these models correctly predict features of the ejection dynamics. We identified ten studies (38-47) that make predictions that can be directly compared to our findings, summarized in Supplementary Tables S1 and S2. Some of these studies predict features in accord with our findings at a qualitative level (Supplementary Table S2). First, several predict large variations in the dynamics between individual ejection events and link this to variability in initial DNA conformations (39,40,45-47). Second, three studies predict an initially increasing velocity (38,40,41). Third, several predict stochastic pausing (38-41,45-47). The nature of the stochastic pausing we observe is much closer in character to that predicted by these simulation studies than the pausing observed in the phage T5 studies with dye labeled DNA, where pauses were mainly observed at specific capsid filling levels.

However, at a quantitative level we do not find good agreement with most of the predictions (Supplementary Table S2). First, almost all the simulation studies predict ejection velocities orders of magnitude higher than we measure, even after considering predicted scaling laws for the dependence of ejection timescales on DNA length/thickness, initial packing density, and viscosity (see Supplementary text for a discussion). Second, although three studies predict initially increasing velocity, none predict an exponentially increasing trend with decreasing packing density like that we observe. Third, although several studies predict pausing, detailed pausing statistics were not reported and predicted pause durations are generally much shorter than we observe (both on an absolute scale and relative to overall ejection time; Supplementary Table S2). Comparing across all the simulation studies, we do not observe any clear trend suggesting that particular simulation methods or model assumptions result in better agreement with our results (e.g., stochastic rotation dynamics vs. Langevin dynamics, different interaction potentials, different initial packing densities and DNA conformations, etc; Supplementary Tables S1 and S2. One simulation study, by Matsuyama and Yano (41), does predict both an initially increasing velocity and velocity magnitudes on the same order as those we measure after applying predicted scaling laws (for this comparison the model parameters required significant extrapolation to match those of phi29) (Supplementary Table S2). However, this study assumes an intermediate-distance attractive interaction between monomers, which is inconsistent with experimental studies of DNA-DNA interaction forces in our ionic conditions (60,64). The ejection force was also predicted to be independent of packing density, which is inconsistent with experimental studies of phages phi29 and lambda (11,24,26,27).

Various factors likely contribute to the disagreement between simulation studies and our experimental findings. Many studies modeled DNA lengths/capsid sizes much smaller than phi29 and some modeled polymer properties different from those of DNA (Supplementary Table S1). The predicted scaling laws used to extrapolate these predictions for comparison have not yet been experimentally verified. More importantly, our evidence for DNA-DNA sliding friction suggests that coarse-grained simulation studies likely would not accurately model such friction since they do not model specific nanoscale structural features and electrostatic interactions of DNA. Since we find that the initial exit velocity is exponentially dependent on packing density, another issue is that most simulation studies assumed lower initial packing densities than that inferred by the latest X-ray studies on phage phi29 (Supplementary Table S1) (68). Moreover, Mahalik *et al*. simulated DNA packaging prior to ejection using driving forces ∼2 fold larger than the experimentally determined internal force in phi29, yet predicted a ∼2-fold lower maximum packing density (47). This raises a question of whether packing densities in coarse-grained simulation models can be accurately related to those in experiments.

Features of the pausing we observe lead us to propose that it is related to a phenomenon in soft matter physics termed “clogging” (72-77), which occurs when non-equilibrium materials such as colloid solutions or granular materials are forced to flow through a constriction. Clogging events that temporarily arrest the flow occur due to spontaneous formation of metastable arrangements of particles. One example is in gravity-driven flow of agitated granular material through a hopper, where particles spontaneously form arch-like structures that span the constriction and block the flow until external agitation breaks them apart (72-74). Another example is in flowing colloid solutions, where Brownian fluctuations play a role in driving clogging/unclogging events (75,76). Clogging events have varying durations and occur over a wide range of particle densities for a given set of system parameters (e.g., particle and constriction size ratio, agitation intensity, driving force, etc.).

Here, we envision that the “particles” that clog are short DNA segments that approximately behave as independent particles on scales much smaller than the persistence length (P≅150 bp). The phi29 genome has ∼130 persistence lengths, so segments separated by >>P behave, to a degree, as independent particles. Studies of clogging also find that its probability increases when there is greater friction between the particles (74,77). Thus, the DNA-DNA sliding friction we find evidence for would promote clogging. The ubiquitous predictions of pausing in simulations of DNA ejection, discussed above, supports the hypothesis that these pauses are related to fluctuations in the local DNA conformation during ejection (38-41,45-47).

Experimental and theoretical studies suggest that clogging across many types of soft matter systems exhibits universal statistical properties. Most notably, the amounts of material that flow between clogging events follow an exponential distribution, consistent with onset being a random event, while the distributions of clog durations have power-law tails (73,78-80,82,83). The pauses we measure show similar features. First, the distribution of lengths of DNA that exit between pauses at high filling follows an exponential distribution (Figure 4A). Second, the distribution of pause durations (Figure 4B) is not exponential but has a power law tail. Third, the fitted power law exponent of 2.3 falls in the range of those measured for other soft matter systems that exhibit clogging (72,73,78-80). Fourth, decreasing the pulling force or increasing the ionic screening decreases the average lifetime of pauses, which is consistent with studies showing that clogging events decrease in duration when the driving force is increased (78,81). Fifth, the pausing frequency increases at high filling levels, consistent with studies showing that the probability of clogging increases with increasing particle density (83,84).

## Conclusions

In summary, we introduce a method for probing the mobility of the tightly packed DNA in viral capsids and shed light on the nature of the friction forces that limit the speed of DNA exit. The initial exit velocity for phi29 is much lower than has been reported for other phages using dye-labeled DNA, and much lower than has been predicted by theory/simulation studies. We provide evidence that the kinetics of ejection are dominated by DNA-DNA sliding friction during the initial 20% of DNA exit and by stochastic pauses that are different in character than those observed previously with phage T5 and different than predicted in simulation studies. We present evidence suggesting that the pausing is connected to the phenomenon of “clogging” observed in nonequilibrium soft matter systems.

## Acknowledgments

We thank Damian delToro for work on setting up the optical tweezers instrument.

## Author contributions

M.F., N.K., and D.E.S. designed research, M.F., N.K., and P.J.J. performed research, M.F., N.K., and D.E.S. analyzed data, P.J.J. edited the manuscript, and M.F. and D.E.S. wrote the manuscript.

