## Supplementary Data for "Role of DNA-DNA sliding friction and non-equilibrium dynamics in viral genome ejection and packaging"

#### Hydrodynamic drag during phi29 ejection

In the model proposed by Grosberg and Gabashvili (1), the hydrodynamic drag force experienced by the ejecting DNA is given by

$$F = \frac{2\eta\pi v L_{DNA}}{\log\left(\frac{d_s}{R_{DNA}} - 1\right)},$$

where $\eta$ is the solvent viscosity, $v$ is the ejection velocity, $L_{DNA}$is the length of the confined DNA, $d_s$ is the interaxial spacing of the confined DNA, and $R_{DNA}$ is the radius of DNA (1 nm). Using the viscosity of water at 23°C, the length of the phi29 genome (19,282 bp), our measured DNA exit velocity at 95-100% filling (~200 bp/s), and an estimated interaxial spacing of 2.48 nm (47) gives a drag force of ~0.01 pN.

#### Theoretical studies of viral DNA ejection

We surveyed the literature for theoretical studies of viral DNA ejection. The discussion in the main text, and summarized in Table S1 and S2, focused on those that can be directly compared with our findings. Other studies among the 48 mentioned in the text are mainly focused on the energetics, mechanics, and thermodynamics of DNA ejection and/or predicting free energies or scaling relationships (6,12,14,16-30,32-38,40-42,44-46). Some studies also considered the effect of knots in DNA (31,39,43).

#### Comparison with simulation studies

The physical parameters associated with phi29 DNA exit and the parameters used in coarse-grained molecular dynamics simulation of bead-spring polymer ejection (2-10, 46) are compared in Table S1. These parameters include the dynamic simulation method (algorithm) used, terms used in the potentials, whether hydrodynamic interactions are included, capsid shape, the monomer size $\sigma$, the initial volume fraction $\phi_0$, the persistence length $\ell_p$, the smallest dimension of the capsid $R_C$ (the radius of spherical capsids and the semi-minor axis for elongated capsids), the length of the chain $L_{chain}$, and the "initialization method" by which the initial conformation of the packed DNA was set. The initial volume fraction for simulation studies was either given in the text or calculated as $\frac{4\pi N}{3}\left(\frac{\sigma}{2}\right)^3/V_c$, where $V_c$ is the capsid volume. The initialization methods used were either packaging by applying a constant force at the pore, meant to simulate viral DNA packaging, or a "random" configuration with some imposed waiting time to allow for chain relaxation before ejection. In studies where multiple values of bending rigidity were investigated, we compared with results from simulations using a persistence length that produced an initial driving force closest to the internal force in phi29. In Ali *et al.* (4), multiple values for the strength of ionic screening of electrostatic interactions were investigated, so we present results from simulations with a Debye screening length of 1 nm, which most closely matches our experimental conditions.

Various simulation studies use parameters, such as DNA length and viral capsid size, that differ somewhat from those appropriate for phage phi29 (Table S1). To compare our measurements with simulation predictions more quantitatively, we also extrapolated the predictions using predicted scaling laws. Several studies that have examined the dependence of ejection time on the system parameters are not in exact agreement regarding the scaling exponents (although they are generally not wildly different) (7,8,15,20,30). We therefore present a range of predicted scaled velocity values considering the range of scaling exponents on initial chain length and initial volume fraction reported in different published studies. The general predicted form of the scaling dependence of ejection time in the pressure-driven ejection regime on the system parameters is $t{\sim}\tau_0\phi_0^{-a}N_0^b$, where $\tau_0{\sim}\eta\sigma^3$ and $\eta$ is the viscosity. The predicted exponents *a* and *b* range from 1.25 to 1.7 and 1.2 to 1.7, respectively (7,8,15,20,30).

We compare the predicted magnitude of the ejection velocity and dependence on volume fraction from simulation studies and analytical models to our measurements in Supplementary Table S2, as well as the occurrence and duration of pauses. In particular, we compare the velocity during the first 10% of the ejection (in the regime where we find the velocity is sharply increasing), $v_{10\%}$, calculated as the average ejection velocity as the volume fraction decreases from $\phi_0$ to $0.9\phi_0$ , where $\phi_0$ is the volume fraction at the beginning of ejection. For analytic models, this was calculated with the model presented in the study using parameters corresponding to phi29. For simulation studies, this velocity was scaled, using the predicted scaling laws mentioned above, to account for the length, volume fraction, viscosity, and monomer size for DNA packaged in phi29. The absolute pause duration $t_p$ was estimated for pauses shown in simulated ejection trajectories occurring during the beginning stages of ejection, where the internal force driving ejection is large (during the first 30% decrease in volume fraction). Since most of the simulations find ejection velocities much higher than we measure, we also compared pause duration relative to ejection timescale, calculated as $t_p$ divided by the time to eject the first half of the chain. Mappings between simulation units and physical units for length and time were given in (2-5,9,10,46) and used to convert $\ell_p$, $R_C$, $L_{chain}$, $v$, and $t_p$ to physical units for comparison with our results.

### Supplemental Figures and Tables

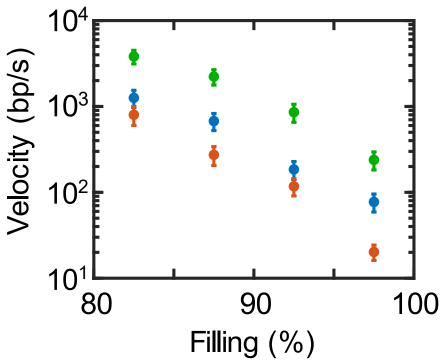


**Figure S1:** Average DNA exit velocity including pauses versus capsid filling level in the high-filling regime in the with 5 pN applied force in the $Na^+$ screening condition (blue, n = 39 events) and $Mg^{2+}$ screening condition (red, n = 31) and with 20 pN applied force in the $Na^+$ screening condition (green, n = 42).

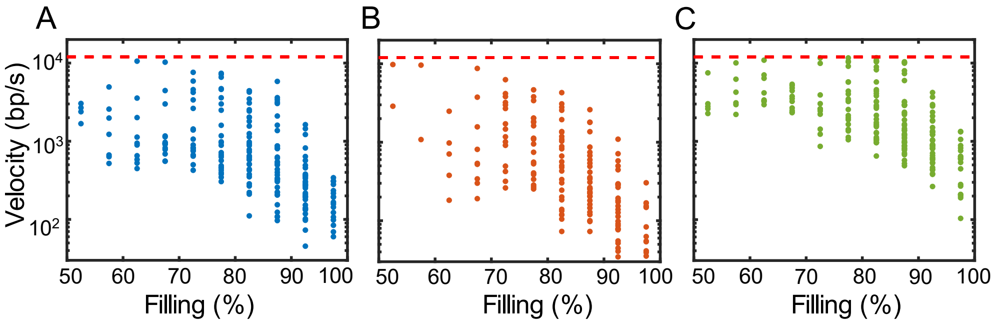


**Figure S2:** DNA exit velocities determined for individual complexes in 5% capsid filling level bins vs. filling level with (A) 5 pN applied force in the $Na^+$ screening condition (blue, n=39 events), (B) 5 pN applied force in the $Mg^{2+}$ screening condition (red, n=31), and (C) 20 pN applied force in the $Na^+$ screening condition (green, n=42). The upper bound for measurable DNA exit velocity is indicated by the dashed red line.

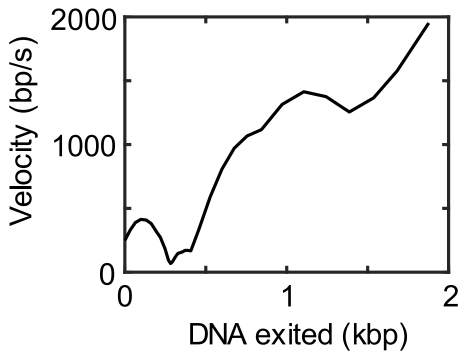


**Figure S3:** DNA exit velocity vs amount of DNA remaining in the capsid plotted for an individual complex in the $Na^+$ screening condition with a 5 pN pulling force. Velocity was calculated using a 0.5 s sliding window applied to the tether length vs. time data after pauses were identified and removed (as described in *Methods*).

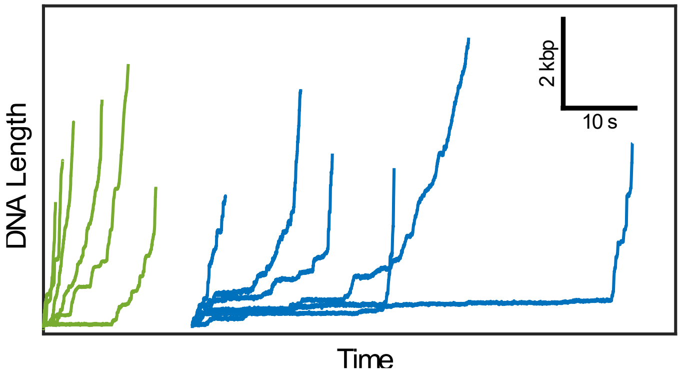


**Figure S4:** Measurements of length of DNA that has exited the capsid vs. time in the $Na^+$ screening condition with 5 pN applied force (blue, n=39 events) or 20 pN (green, n=42).

| Reference | Method | Potential | Hydrodnamic interactions | Capsid shape | $\sigma$ (nm) | $\phi_0$ | $\ell_p$ (nm) | $R_c$ (nm) | $L_{chain}$ (nm) | Initialization method |
|---|---|---|---|---|---|---|---|---|---|---|
| Phi29 (current study) | | | Yes | Elongated | 2 | 0.65 | 50 | 16 | 6555 | Packaging |
| Ali *et al.* (2006) | SRD | LJ†/FENE/KP | Yes | Elongated | 2.5 | 0.45 | 25 | 6.5 | 250 | Random |
| Ali *et al.* (2008) | SRD | WCA/FENE/KP | Yes | Spherical | 2.5 | 0.45 | 25 | 7.5 | 250 | Random |
| Ali *et al.* (2011) | SRD | WCA/FENE/KP/DH | Yes | Spherical | 2.5 | 0.45 | 25 | 7.5 | 250 | Random |
| Matsuyama *et al.* (2012) | LD | LJ/Hookean/KP | No | Spherical | 1 | 0.46 | 13 | 3.0 | 100 | Random |
| Marrenduzzo *et al.* (2013) | KJD | WCA/FENE/KP/DH/CL | No | Spherical | 2.5 | 0.11 | 50 | 22.5 | 1600 | Packaging |
| Mahalik *et al.* (2013) | LD | WCA/Hookean/KP | No | Elongated | 2.5 | 0.37 | 60 | 16 | 3500 | Packaging |
| Piili *et al.* (2015) | SRD | WCA/FENE | No | Spherical | 8 | 0.63 | 4 | 27 | 1600 | Packaging |
| Linna *et al.* (2017) | SRD | WCA/FENE/KP | Yes | Spherical | 2 | 0.43 | 22 | 7.7 | 400 | Packaging |
| Huang and Hsiao (2019) | LD | WCA/Hookean | No | Spherical | N.R. | 0.4 | N.R. | N.R. | N.R. | Packaging |
| Park and Sung (2021) | LD | WCA/FENE/KP | No | Spherical | 2.5 | 0.31 | 50 | 16 | 1650 | Packaging |

† Attractive part of the Lennard-Jones term was excluded.

**Table S1**: Summary of methods and parameters used in bead-spring polymer simulation models of DNA ejection compared with experimental properties of DNA and phage phi29. $\phi_0$ refers to the initial DNA packing volume fraction at the start of ejection. $\ell_p$ refers to the persistence length of the polymer. $Lchain$ refers to simulated DNA length. σ refers to the polymer thickness (diameter of the monomers). $Rc$ refers to the minimum dimension of the capsid (equal to the radius for spherical capsids and the semi-minor axis for elongated capsids). "Initialization method" refers to how the initial DNA conformation was determined; "packaging" refers to studies where a simulation of DNA packaging was done prior to simulations of ejection. "Method" refers to the simulation method used: Stochastic Rotation Dynamics (SRD) or Langevin Dynamics (LD). "Potential" refers to the polymer bead-bead interaction potential used in simulations. Different terms in the potentials include: a Lennard-Jones (LJ) term or a Weeks-Chandler-Andersen (WCA) term to model excluded volume interactions, a finite extensible nonlinear spring (FENE) or Hookean spring term, a Katky-Porod (KP) bending rigidity term, a Debye-Huckel (DH) electrostatic interaction term, and a "Cholesteric" term (CL), which introduces interstrand angular correlations as observed in some DNA liquid-crystal phases.

| Reference | $v_{10\%}(\phi)$ direction | $v_{10\%}(\phi)$ form | $v_{10\%}$ (nm/s) | Scaled $v_{10\%}$ range (nm/s) | Pauses observed | Absolute $t_p$ (s) | Relative $t_p$ |
|---|---|---|---|---|---|---|---|
| Phi29 (current study) | Increasing | Exponential | $9 \times 10^1$ | $9 \times 10^1$ | Yes | 2 | 0.3 |
| Ali *et al.* (2006) | Increasing | Linear | $3 \times 10^8$ | $(5 \times 10^8, 3 \times 10^9)$ | Yes | $^\dagger$ $8 \times 10^{-7}$ | 0.1 |
| Ali *et al.* (2008) | Decreasing | Linear | $^\dagger$ $1 \times 10^7$ | $^\dagger$ $(2 \times 10^7, 1 \times 10^8)$ | Yes | $2 \times 10^{-7}$ | 0.04 |
| Ali *et al.* (2011) | Increasing | Linear | $1 \times 10^8$ | $(1 \times 10^8, 9 \times 10^8)$ | Yes | N.R. | N.R. |
| Matsuyama *et al.* (2012) | Increasing | Linear | $4 \times 10^3$ | $(1 \times 10^2, 9 \times 10^2)$ | Yes | N.R. | N.R. |
| Marrenduzzo *et al.* (2013) | N.R. | N.R. | $^{\dagger\ \S}$ $3 \times 10^5$ | $^{\dagger\ \ddagger}$ $(2 \times 10^6, 9 \times 10^6)$ | Yes | N.R. | N.R. |
| Mahalik *et al.* (2013) | N.R. | N.R. | $^\dagger$ $4 \times 10^7$ | $^\dagger$ $(3 \times 10^2, 6 \times 10^2)$ | Yes | $^\dagger$ $2 \times 10^{-6}$ | 0.05 |
| Piili *et al.* (2015) | Decreasing | Exponential | N.R. | N.R. | N.R. | N.R. | N.R. |
| Linna *et al.* (2017) | Decreasing | Exponential | N.R. | N.R. | N.R. | N.R. | N.R. |
| Huang and Hsiao (2019) | Decreasing | Power law | N.R. | N.R. | N.R. | N.R. | N.R. |
| Park and Sung (2021) | N.R. | N.R. | $^\S$ $6 \times 10^8$ | $(2 \times 10^9, 3 \times 10^9)$ | Yes | $^\dagger$ $5 \times 10^{-7}$ | 0.2 |
| Gabashvili and Grosberg (1992) | Decreasing | Exponential | $3 \times 10^5$ | N/A | N/A | N/A | N/A |
| Inamdar *et al.* (2006) | Decreasing | Exponential | $2 \times 10^3$ | N/A | N/A | N/A | N/A |
| Sakaue and Yoshinaga (2009) | Decreasing | Power law | $1 \times 10^{10}$ | N/A | N/A | N/A | N/A |
| Wang *et al.* (2012) | Increasing | Linear | $1 \times 10^4$ | N/A | N/A | N/A | N/A |

$^\dagger$ Value estimated from plot of ejection trajectories or average ejection trajectory.
$^\ddagger$ Viscosity of fluid assumed to be that of water at 300K.
$^\S$ Average ejection velocity - initial velocity not reported.

**Table S2**: Comparison of our measured findings on DNA exit from phi29 with predictions of various simulation studies (blue) and analytic studies (orange). Ejection velocities were calculated and scaled to account for differences in model parameters as described in the Supplementary text. Pause duration $t_p$ was measured for pauses shown in individual simulated ejection trajectories (range during which $\phi$ decreased by 30%) and adjusted to a relative value by dividing the time to eject the first half of the polymer. Values that were not reported are listed as *N.R*. See the section of the Supplementary text titled "Comparison with simulation studies" for further details.
